# Prediction and experimental evidence of different growth phases of the *Podospora anserina* hyphal network

**DOI:** 10.1101/2023.02.08.527651

**Authors:** Clara Ledoux, Florence Chapeland-Leclerc, Gwenaël Ruprich-Robert, Cécilia Bobée, Christophe Lalanne, Éric Herbert, Pascal David

## Abstract

The growth of the network of a filamentous fungus is monotonous, showing an ever increasing complexity with time. The components of network growth are very simple and based on two mechanisms, the elongation of each hyphae and their multiplication by successive branching. These two mechanisms are sufficient to produce a complex network, and could only be localized at the tips of the hyphae. However, branching can be of two types, apical or lateral, depending on its location on the hyphae, imposing the redistribution of the necessary material in the whole network. From an evolutionary point of view, maintaining different branching processes, with its associated cost, is intriguing. We propose in this work to discuss the advantages of each of the two branching types using a new observable for the network growth allowing to compare growth configurations. For this purpose, we build on experimental observations of the *Podospora anserina* thallus growth, allowing us to feed and constrain a lattice-free modeling of this network based on a binary tree. First, we report the set of statistics related to the branches of *P. anserina* that we have implemented in the model. Then, we build the density observable, allowing us to discuss the growth phases. We predict that the evolution of the density is not monotonic but presents a marked stationary phase, clearly separating two other phases of growth. The location of this stable region appears to be driven solely by the growth rate. Finally, we show that the density is a good observable for stress differentiation.

## 1 Introduction

The achievement of filamentous fungi in colonizing terrestrial ecosystems can be largely attributed to their flexible morphology, and more specifically to their ability to form an interconnected hyphal network, the mycelium, based upon some fundamental cellular processes, such as hyphal tip growth, septation, hyphal orientation, branching and fusion (also known as anastomosis).^1^ Hyphal branching has been well described and appears to both increase the surface area of the colony, which enhances nutrient assimilation, and mediate hyphal fusion events that are important for exchange of nutrients and signals within the same mycelium.^2^ Therefore, the architecture of the fungal network is clearly not fixed, but must continually adapt to local nutritional or environmental cues, damage or fungivore attacks. Such a network is constituted of apical branches (or leading hyphae), which are the first to invade new territory and are generally engaged in nutrient acquisition and sensing of the local environment, whereas behind the colony edge, subapical cells generate new hyphae by lateral branching.^3^ An earlier description of such fungal network states that hyphal tips at the biomass edge are those associated with exploration while hyphal tips behind the biomass front are most associated with resource exploitation.^4^

All living systems are *a priori* motivated by access to resources and reproduction and constrained by the environment and its internal metabolism. Evolution proceeds from the selection of random mutations in the genotype that will retroact on the phenotype. This is the genotype-phenotype relationship as described in^5, 6, 7^ for different biological systems. This process seems to give rise to simple systems in the sense of Kolmogorov.^8^ This can be understood as a bias towards simplicity. Based on the algorithmic description of Kolmogorov,s complexity,^9^ *i.e*. the size of the smallest program necessary to generate information, it is found that a short program is more likely to appear more frequently. Based on this principle, it was recently suggested^10^ that simplicity and symmetry could emerge spontaneously from the evolutionary process. The growth of a branching network can be most simply obtained by placing an active body at each tip, the Spitzenkörper or SPK in the case of ascomycete filamentous fungi. This structure is belived to regulate the delivery of cell wall-building vesicles to the apical cell surface, thereby allowing the elongating of hyphae, as well as the generation of a new tip, i.e. a branching event.^11, 12, 13^ However, the branching process is not only observed at the apexes, where apical branching occurs, but also in the subapical regions distant from the tip, virtually anywhere on the network. The latter is named lateral branching. This imposes the redistribution of the growth machinery and the resources within the network, which mechanically increases its complexity. The question that arises concerns the interest of relying on both branching types for the growth of the fungal network, so that they have been selected and kept.

*Podospora anserina*^14^ is a coprophilous filamentous ascomycete, a large group of saprotrophic fungi that mostly grows on herbivorous animal dungs and plays an essential role within this complex biotope in decomposing and recycling nutrients from animal feces. *P. anserina* has long been used as an efficient laboratory model to study various biological phenomena, especially because it rapidly grows on standard culture medium, it accomplishes its complete life cycle in only one week, leading to the production of ascospores, and it is easily usable in molecular genetics, cellular biology and cytology.

As was already discussed in,^15^ the growth of hyphal network expansion and structure of *P. anserina*, were characterized under controlled conditions. Temporal series of centimetric image size of the network dynamics, starting from germinating ascospores, were produced with a typical micrometric resolution. The image reconstruction steps were completely automated and allowed easy post-processing and quantitative analysis of the dynamics. Based upon the two main processes that drive the growth pattern of a fungal network, *i.e*. apical growth and hyphal branching, we have proposed^16^ a two-dimensional simulation based on a binary-tree model, allowing us to extract the main characteristics of a generic thallus growth. In particular, we showed that, in a homogeneous environment, the fungal growth can be optimized for both exploration and exploitation of its surroundings with a specific angular distribution of apical branching. This numerical experience is obviously far from the *in vivo* growth of a saprophytic fungus such as *P. anserina*. However, it constitutes an excellent starting point to describe the thallus growth using mathematical concepts and language. Indeed, the objective of our mathematical modelling is to reduce a complex biological system to a simpler model, which is able to partly reproduce, or even better predict, the real system. A recurrent question is then to find the optimal degree of simplification for such a model, which should be neither too simple to avoid straying from realistic predictions, nor too complex to solve using numerical methods.^17^

The structure of this article is as follows. In the *Results* section, we first recall the methodology used to model the thallus, then we describe the statistics extracted from observations that were integrated into the simulation. These concern the lateral and apical branches, the curvature and the symmetry of the growth. Finally, we describe the density observable and derive predictions. In the *Discussion* section, we compare with direct observations made on the thallus growth in two configurations, the first on a standard culture medium, and the second on a depleted medium.

## 2 Results

### 2.1 Modelling the thallus growth

Direct observation of the growth of the fungus network shows a monotonous, ever increasing complexity. This is particularly obvious via observables such as the number of branches, or the total length of the network. This remains true, even when the culture medium is modified, with for example a quantity of nutrient depleted.^16^ In the latter case, we found that the growth rate is affected but not the growth itself, *i.e*. only the value of the exponential growth parameter is affected. However, these observables are global aggregates and do not capture finer effects such as a change in the spatial distribution of matter. In this section we introduce a new observable that combines both the amount of matter and its distribution via the density of the network.

In a previous article^16^ we discussed the foundations of a model built to describe the growth dynamics of *P. anserina* branching network. In order to ease the reading of the work presented in the following, we recall here its main characteristics.

The observed network is composed of interconnected branches, called hyphae, whose ends are the apexes. Growth in hyphal length is achieved by adding material to the apex. The connections correspond to branches that may appear at the apex—called apical branching—or along a hypha—called lateral branching.

The simulation of network growth is based on the reproduction of these basic elements. The apical branching can only show one additional apex. We therefore rely on a binary tree to which further allows lateral branching events.

In the nomenclature used, *V*_1_ are the tips (or apexes) of the branches, *V*_3_ are the vertices corresponding to the connection between three branches, *V*_1*ℓ*_ are the apexes of the lateral branches and *V*_3*ℓ*_ are the nodes of these branches. In addition to these biologically defined objects, we have taken care to distinguish the crossings (overlaps) of hyphae from real vertices. These geometric intersections, called *V*_3*i*_ in the following, should not be confused with anastomosis (hyphae merging). This distinction is necessary to compare with observations from the experimental conditions. During growth, the network is arranged on a surface, but is not constrained in its upper part. Thus, overlaps can occur when two hyphae are brought to cross each other, which makes the frequency of overlap between hyphae important. In addition, the acquisition technique, based on a light intensity contrast between the hyphae and the background, does not allow to discriminate an overlap from a fusion.

The simulation makes it possible to distinguish *V*_3_ and *V*_3*l*_ from geometric vertices. This makes it possible to correct the magnitudes relating to the vertices *V*_3_ and *V*_3*l*_ that were observed experimentally. We check the relevance of the simulation by comparing the ratio of the vertices labeled 1-body (*V*_1_ and *V*_1*l*_) to the other vertices of the image, and this as a function of time, for both the experimental data and the simulation.

In the simulation a set of random variables are used to fix the values of the growth of each apex and the branch angles. We give in^16^ the details of the probability laws used as well as their parameters which were, for some of them, obtained by the analysis of the experimental data.

#### 2.1.1 Experimental observations implemented in simulation

In this section, we describe the observations implemented in the modeling following our previous work. In particular, we relied on statistical descriptions of the branches to discuss their spatial and temporal distribution.

##### Apical branching dynamics

In order to implement the apical branching statistics, we estimated the distribution of the distance *L* between two successive apical branches (*V*_3_). The distribution and corresponding cumulative law are shown in figure 1A for experimental data. Branching statistical behavior is separated into two distinct phases: a first phase of latency, or apical dominance well described in the literature, see for example,^2,18^ and a second phase, which is well recovered by a memoryless law. The apical dominance region was estimated to *L*_0_ = 180 ± 30*μ*m and the value of the rate of the exponential distribution to *α* = (10.4±3.9). 10^−3^*μ*m^−1^. The apical branching is not located exactly at the apex but slightly behind it. This behaviour is named subapical branching in the literature.^19,20^ In this work the expressions *apical branching* and *subapical branching* refer to the same process. The distance *L*_api_ corresponding to the length between the branch and the apex was also recorded. The corresponding mean and standard deviation of the distribution (not shown) were found to be 41 ±11 μm and they are shown with the green line in figure 1B.

**Figure 1.**
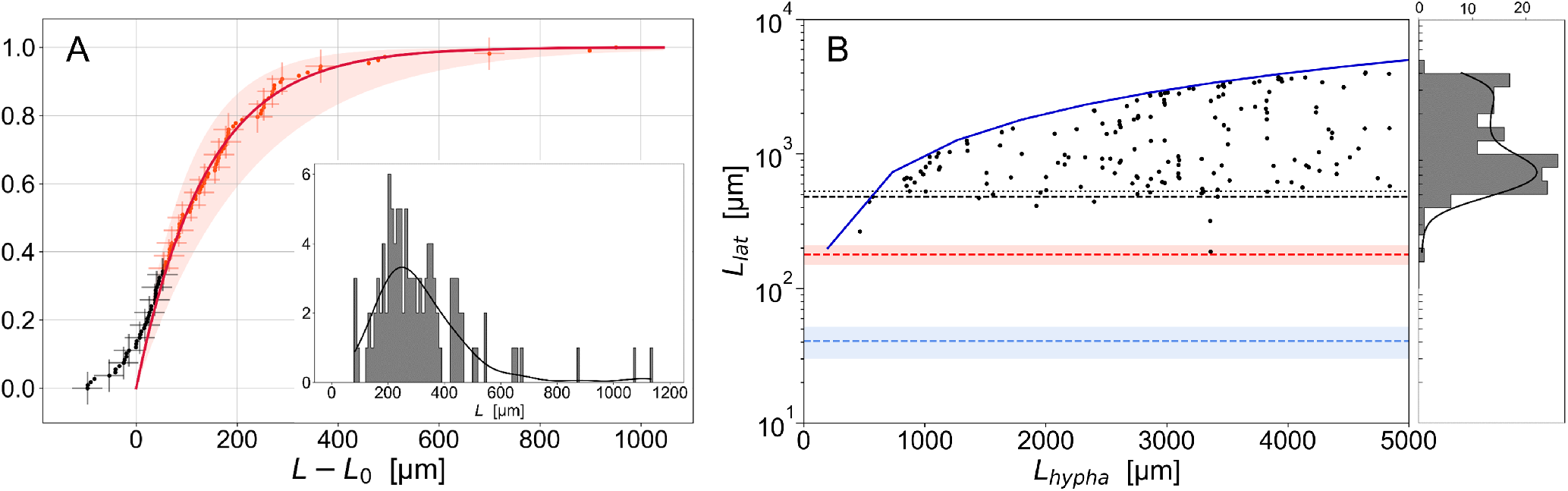
(A) Cumulative law of the distribution (in inset, bins 10*μ*m) of *N* = 109 lengths between two consecutive apical vertices *V*_3_. The black and red markers transition is defined by the maximum slope, found at 230 ± 5 μm from the apex. The solid red line is an exponential fit of the data shown in red with 1 – 2^−α(*L–L*^0^)^. The data were manually shifted by *L*_0_ = 180 μm. Using a diagonal covariance matrix, the exponential fit parameters were found to *α* = (10.4 ± 3.9).10^−3^ μm^−1^, *R*^2^ = 0.99. We made use of *R* squared 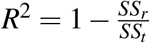, with *SS_r_* the residual sum of squares and *SS_t_* the total sum of squares to discuss the quality of the fit. The red area corresponds to one standard deviation. (B) Lengths *L*_lat_ between the apex and the lateral vertex *V*_3*ℓ*_, measured when the branch appears in function of *L*_hypha_, the length of the hypha when the branch appears. The dark blue solid line corresponds to *L*_lat_ = *L*_hypha_. 95% (resp. 90%) of the data points are above the black dashed line at 480 μm (resp. black dotted line at 530 *μ*m). The red dashed line corresponds to *L*_0_ = 180 *μ*m the apical dominance, as defined in the left figure. The blue dashed line corresponds to the mean and standard deviation of the distribution (not shown) of *L*_api_ = 41 ± 11 *μ*m, defined as the length between the apex and the apical branch at the time of branching (see text for details).

##### Lateral branching dynamics

The dynamics of lateral branching was found to be more subtle. First we discuss the correlation with the distance to the apex. We show in figure 1B the distribution of lengths *L*_lat_ between a lateral branching and the corresponding apex. There is not apparent correlation between a branch and *L*_lat_. However, a region where the probability of observing a lateral branching is extremely low, clearly emerges from the data. We separated the population into two subparts, one composed of 95% of the samples. This is indicated with a black dashed line. We are then allowed to safely conclude that the lateral connections appear more than 480 *μ*m from the apex. This corresponds to the apical dominance behaviour observed for apical branching, but with an associated length about three times higher. In case where the operator has a collection of images of the network with a high enough temporal resolution, it is trivial to distinguish between apical and lateral branches. If, on the other hand, only one image of the network is available, this distinction becomes delicate. Particularly since the branching at the apex is sub-apical, as already discussed. Interestingly, the difference in apical-dominance lengths is a clear parameter for distinction between the two types of branches.

Outside the region of apical dominance, the lateral branches follow their own dynamics. We report in figure 2 the distance Δ*L* between two successive (in time) lateral branches, separated by a duration Δ*t*. We observe that the temporal distribution follows a memoryless exponential distribution, with rate parameter (41 ±19). 10^−3^ min^−1^, *R*^2^ = 0.94. Note that here and in the following, unless otherwise stated, we have used the base-2 exponential function. We made use of 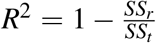, with *SS_r_* the residual sum of squares and *SS_t_* the total sum of squares to discuss the quality of the fit. Note that the laws are linearized to achieve the fit using the logarithm function. Here, *R*^2^ is an indicator of the fit and thus of the logarithm of the law. In the spatial domain, we observe two populations. The observed distribution is compatible with two decorrelated dynamics, which we adjusted based on the mixture of an exponential distributions with a continuous uniform distribution. The procedure is the following. We first fit the data using an exponential distribution Γ_1_2^−γ_1_Δ*L*^ and used the fit parameter *γ*_1_ = (3.3 ±0.4).10^−3^ *μ*m^−1^ (*R*^2^ = 0.80) to proceed with a second fit, which is a combination of the two distributions Γ_1_2^−*γ*_1_Δ*L*^+*r* to determine Γ_1_≈30 ±3 and *r*≈1 ±0.5, *R*^2^ = 0.84. We can then estimate that 22 ±11% of the lateral branch population is driven by the continuous uniform distribution. We can also estimate the associated probability as (0.11 ±0.05).10^−3^/μm/h. We can therefore conclude that lateral branching is driven by two simultaneous behaviors. On the one hand, an intense dynamic near an already existing lateral branch is exponentially distributed. On the other hand, the probability of the emergence of a lateral branch is uniformly distributed. Both populations being numerically of the same order of magnitude.

**Figure 2:**
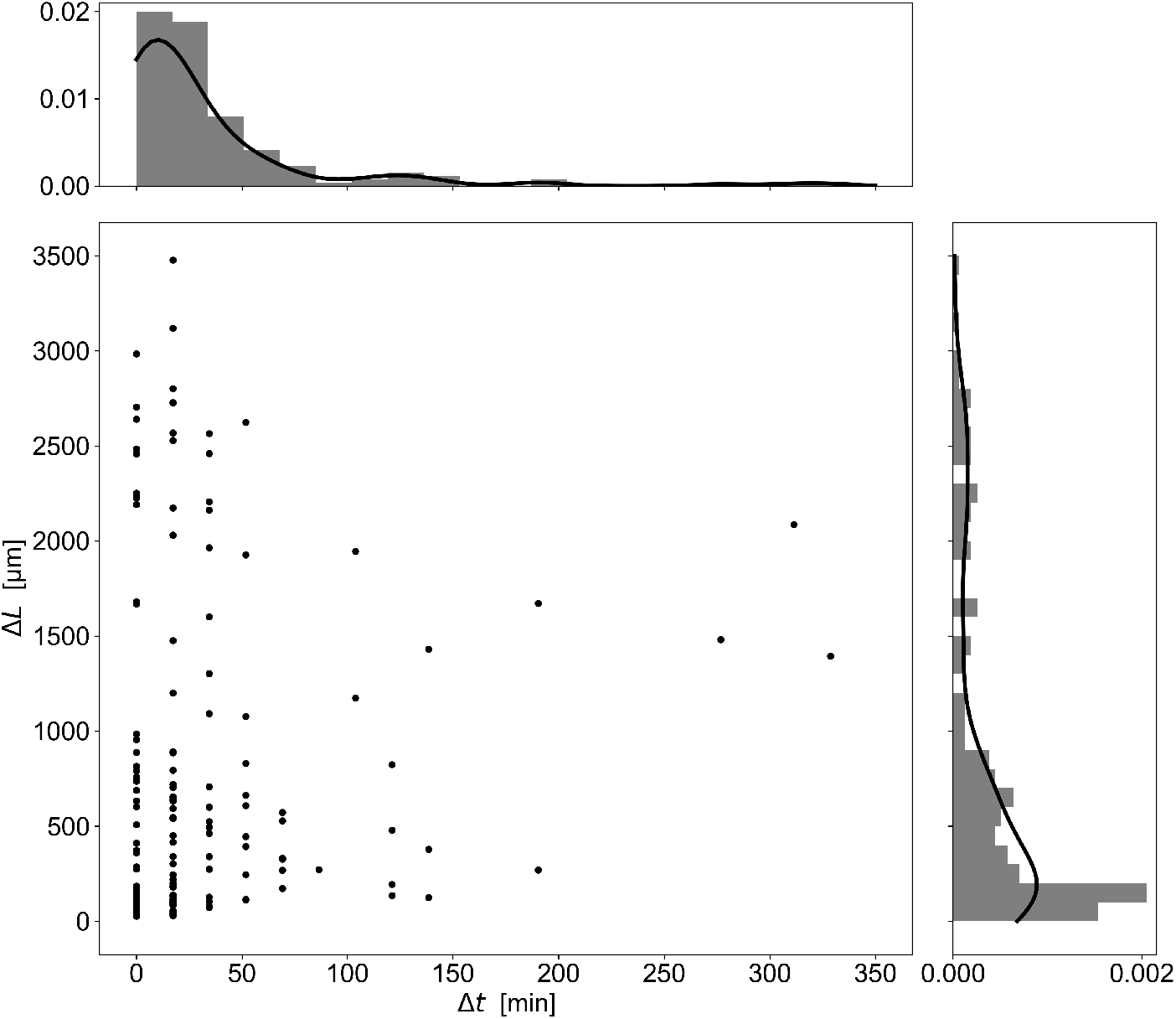
The main figure shows the spatial Δ*L* and temporal Δ*t* distance between two successive lateral branching, taken in chronological order, for *N* = 156 samples. Solid black lines represent the probability density functions associated to the distributions of Δ*L* and Δ*t*.

Finally we found that the scenario of lateral branching dynamics is i)-subject to a region of apical dominance and that ii)-two probability laws are needed to describe the distance between two successive branching, highlighting two different mechanisms. Indeed, two successive branches can appear at a long distance from each other, marking a nucleation process depending on local random fluctuations of resources and building material of hyphae.^21, 22, 23^ The successive branches can also appear close to each other, which is the most likely. This is the signature of an interaction with the environment. Resources absorbed by an apex increase the concentration of materials necessary to branch in the immediate vicinity of the apex. Thus the branches appear by burst in the vicinity of a first branch appearing without predictable location. In order to implement this complex behaviour the nucleation of any lateral branches is determined by the local curvature of the branch during its growth, as is discussed in the following and described Fig 3. Beyond a critical value the position of the step is memorized and a probability law manages the emergence of a branch at each generation.

**Figure 3.**
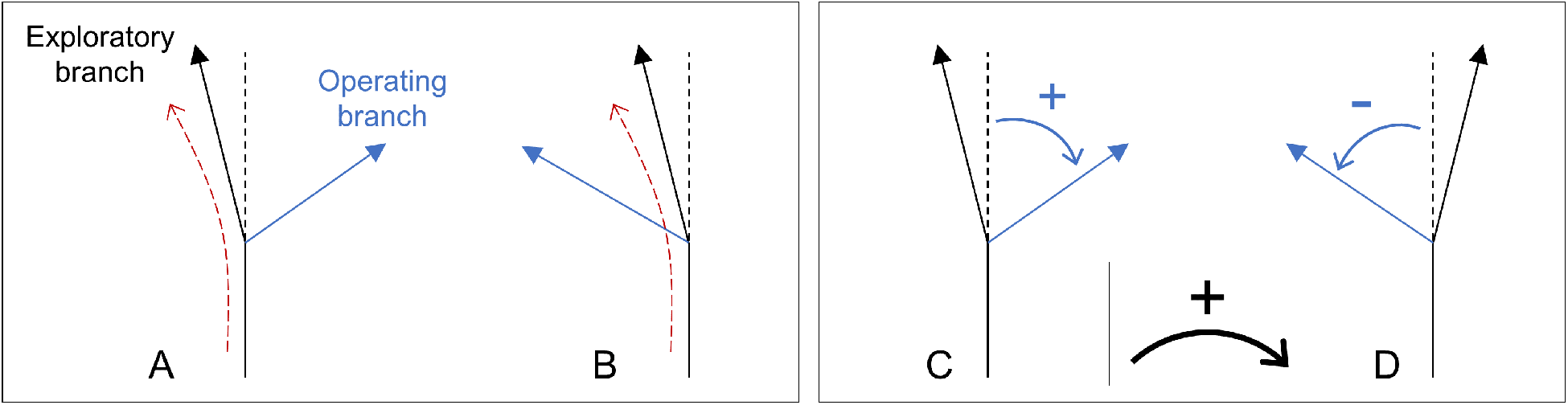
Symmetry of local (left) and global (right) branching. Left, direction of subapical and lateral branching compared to the local curvature of the hypha (subapical branching is shown). (A) is in the opposite direction, (B) is in the same direction. We found 82% and 72% for subapical and lateral branchings respectively corresponding to configuration A. Right, clockwise (C) and anti-clockwise (D) direction of the subapical (large angle apical branching) and lateral branching. We found 54% and 56% for subapical and lateral branchings respectively corresponding to configuration C. All collections are composed of 198 samples. The error is 5% in all cases.

##### Spontaneous curvature

The observation of the trajectory of all the apexes clearly shows, apart from any branching process, that the growth is not rectilinear. We give in this paragraph an estimate of the spontaneous curvature, compared to the rectilinear trajectory, called tortuosity in the following, and the way to take it into account in the simulation. For that purpose, we define “a step of growth”, smaller than the distance between two apical branches. The orientation is determined by a probability law, parameterized according to the previous step. The probability law reads *A* (*θ* – *θ*_0_)^*a*^ (*θ* + *θ*_0_)^*b*^ where *θ*_0_ defines the angular range, *a* > 0 and *b* > 0 are two constant shape parameters, and *A* scales the amplitude of the curvature. This is worth noticing chirality is broken if *a* ≠ *b*. Following the three types of branches described experimentally—*exploratory, operating branches*, and *lateral branches*—we made use of three sets of separate parameters for this propagation mode. Parameters were determined experimentally. It is interesting to note that while the spontaneous curvature of the hypha may be a marker of different spatial occupation strategies, it does not seem to impact the total length of the hypha produced by the network. Thus, we constructed the tortuosity as the arc-chord ratio *L_p_*/*L*_tot_ of the length *L_p_* of the network composed only of nodes of degree 1 and 3, *i.e*. pruned from the curvature, with the total length of the network *L*_tot_. Tortuosity was found in accordance for culture media M2 and M0 (resp. 0.921±0.009 and 0.925±0.008) without any particular correlation in time or space. Although this is not the case in our work on poor and rich culture media, tortuosity can be expected to be different in the case of mutants showing less rectilinear elongation.

##### Branching Chirality

We verified that lateral and apical branches do not spontaneously generate a global symmetry breaking (*i.e*. chirality breaking) by comparing a collection of branches with a defined positive clockwise rotation direction, see Figure 3-right. For apical branches on M2 culture medium, which we define as the reference condition, a binomial test is used to assess whether the frequency of occurrence of positive orientations (measured at 54%) deviates from a theoretical probability of 0.5. The observed p-value was estimated at ≈0.27. Thus, orientation in the positive or negative direction is considered equiprobable. We compared the apical and lateral branches on culture media M2 (56%) and M0 (56% and 58%) to the reference using a Dunnett’s test. The p-values were found to be well above 0.05. Finally we can conclude that probabilities of clockwise and counterclockwise orientation are all found to be consistent with the equiprobability assumption.

We have already discussed the sub-apical rather than apical nature of the apical branching. Combined with the tortuosity (see the paragraph named *Spontaneous curvature*) of the hypha the question arises as to whether the orientation of the operating hypha depends on it, as can be seen in figure 3-left. We have reproduced the previous procedure to test this hypothesis. We found for the binomial test with a probability of success *p* = 0.5 applied to the reference condition a p-value much lower than 0.05 allowing us to clearly reject the equiprobability hypothesis. Surprisingly, the p-value calculated by Dunnett’s test with Bonferroni correction is found to be 2.10^−7^ on the M0 culture medium, which rules out the correlation with the orientation of the mother hypha. Finally, we can retain that the right/left position relative to the parent segment of a lateral and apical branch is only given by the curvature of the parent segment, except in the case of the M0 culture medium for which it is not possible to measure any correlation.

This more complete and realistic simulation allows to take into account some of the more subtle effects observed on the experimental data while introducing a relatively small number of empirical parameters as well as probability laws. As for the previous simulation, time and space parameters were calibrated from the experimental data. Figure 4 allows to compare typical experimental data and simulation for 15 hours of growth duration.

**Figure 4.**
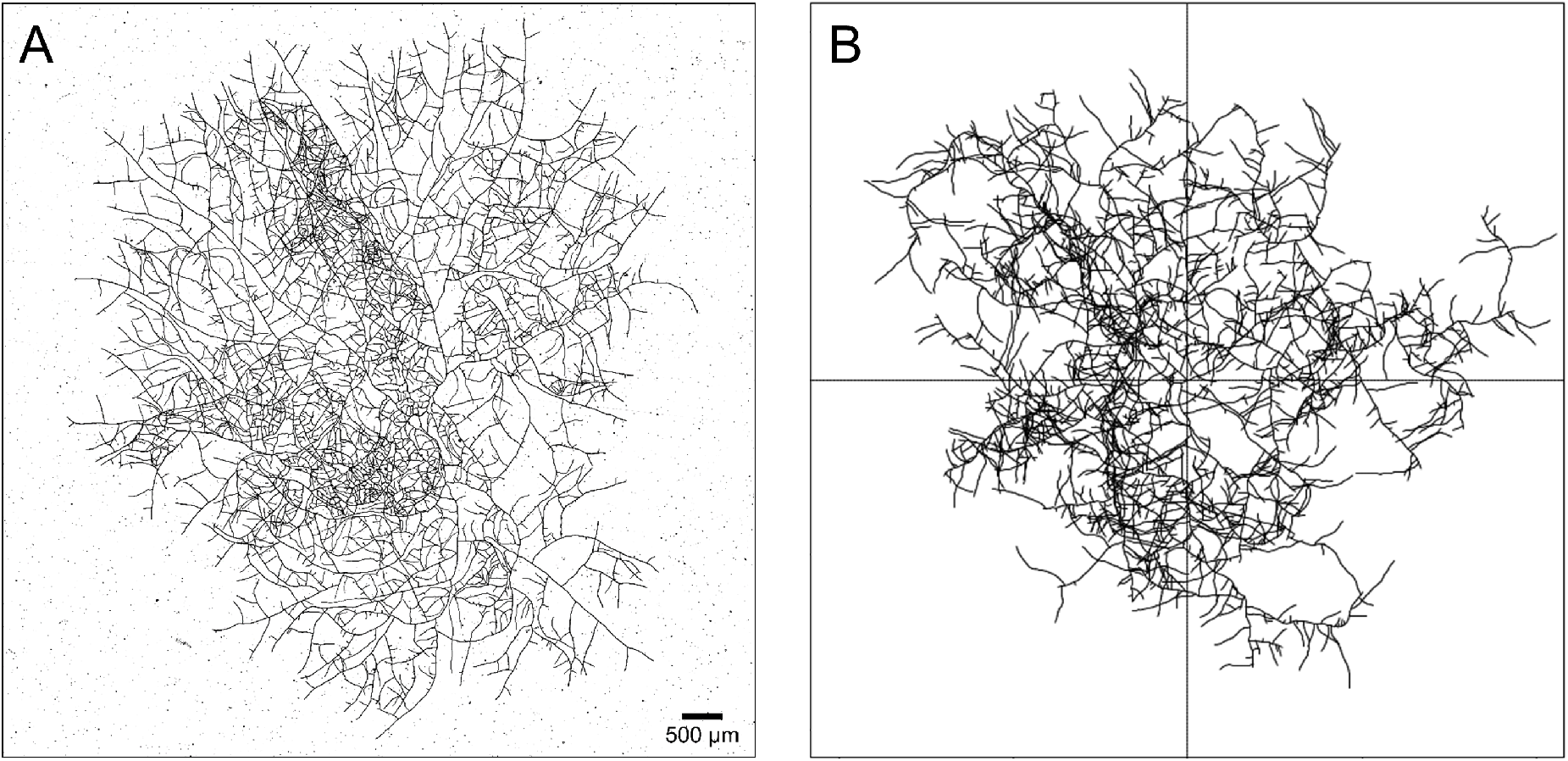
(A) Thallus of *P. anserina* reconstructed from 3 ×4 tiles, extracted from experiment previously discussed in^15^ at *t* = 15 h after the ascospore germination. (B) simulation of the growth of *P. anserina* after the same duration growth, the simulation time were scaled on experimental time.

#### 2.1.2 Different growth phases

The monitoring of the growth of the network can be quantified at each moment thanks to the counting of the vertices of different natures in the network. This work is made simple in the case of the simulation because there is no ambiguity about the qualification of the different nodes: *V*_1_, *V*_1l_, *V*_3_, *V*_3*ℓ*_. In the following, we will rely on the number of *V*_1_+*V*_1*l*_ (apexes) vertices as a function of time to monitor the growth of the thallus. However, this number does not allow the differentiation of the different phases of growth, nor the study of the external constraints that can be applied to this growth. For this, the spatial extension of the thallus must be taken into account. To this end, we define a characteristic observable *density* 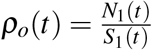 where *N*_1_(*t*) is the number of all 1-body vertices (*N*_*V*_1__ and *N*_*V*_1*l*__) and *S*_1_(*t*) is a characteristic surface generated by the spatial distribution of *V*_1_. We propose to define this surface as the square of the characteristic length generated by the spatial distribution of the *V*_1_ vertices. In a previous work,^16^ we introduced the inertial tensor *I* of the spatial distribution of the vertex *V*_1_. We will briefly recall in the following the derivation method of the characteristic length. Let us first write the expression of this tensor:

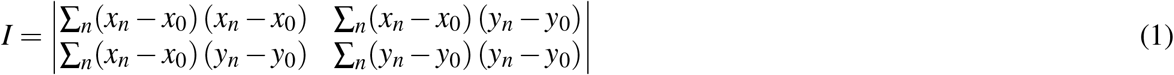

with (*x_n_, y_n_*) the coordinates of the *V*_1_(apical) collection of locations at time *t* and (*x*_0_, *y*_0_) the average of this collection. Diagonalization of this tensor allows to derive two eigenvalues (*λ*_1_ and *λ*_2_) and two eigenvectors, which are the main axes of the *V*_1_ vertices cloud. These eigenvalues correspond to the dispersion of the vertices in the plane and have the dimension of the square of a length. We can then directly derive a surface by calculating the square root of their product 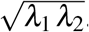. This surface is characteristic of the *V*_1_ distribution for each time step and we propose to use it as a proxy for *S*_1_. We show in figure 5A the temporal variation of the two roots of the eigenvalues, 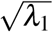 and 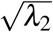, of the *V*_1_ vertices extracted from networks obtained using both a simulation and an experiment.

**Figure 5.**
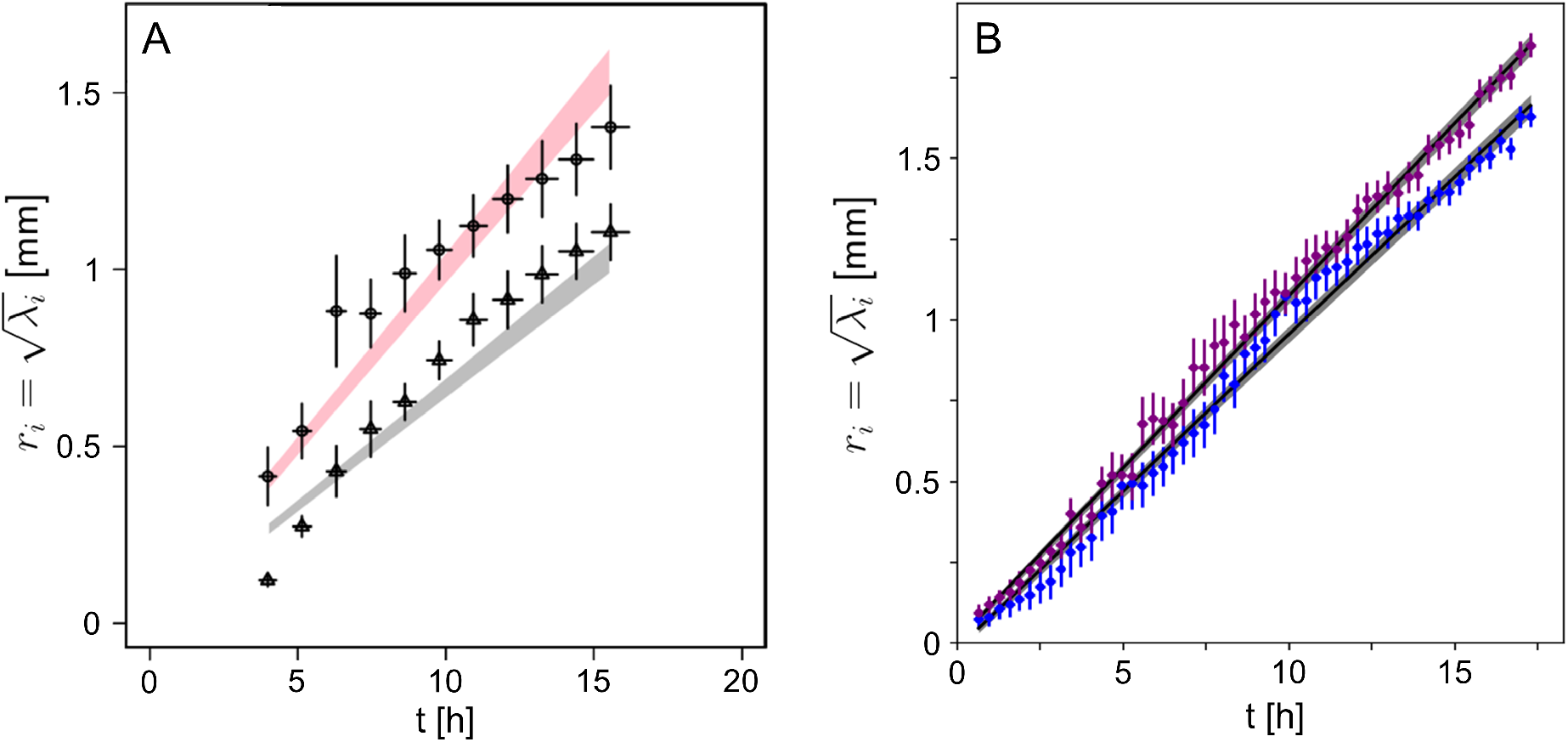
Roots of the eigenvalues 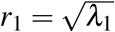 and 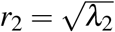 of the apexes (*V*_1_) cloud distribution—see equation 1—as a function of growth time. (A) corresponds to the modelling of the growth, points correspond to the empirical data of simulation, shaded area are the theoretical function *r_i_* = *B_i_t*. (B) corresponds to experimental data, obtained from experiment M2_1_, see text for details. The colored area shows one standard deviation for both fits. Corresponding slopes *B_i_* are 0.098 ± 0.002, 0.107 ± 0.002 mm.h^−1^ with *R*^2^ > 0.99. In this last case the data have been manually shifted by *t*_0_ = 1 h±10 minutes. Corresponding sphericity 2*λ*_2_/(*λ*_1_ + *λ*_2_), with *λ*_1_ ≥ *λ*_2_, is found to be constant at 0.88±0.02.

The spatial extension of the thallus is constrained by its boundary conditions. The surface must be zero for *t* = 0 and must converge asymptotically to a finite value in the long run. These constraints must be reflected in the chosen law to adjust the time behaviour of the eigenvalues. The law which best describes the evolution of the roots of eigenvalues is 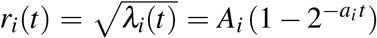 where *i* refers to the two eigenvalues *λ*_1_ and *λ*_2_, and *a_i_, A_i_* are positive constants. Apart from the spatial extension, both longitudinal and transverse growth velocity of the *V*_1_ distribution can be extracted by deriving *r_i_* with respect to time, *i.e*. *v_i_*(*t*) = *∂_t_r_i_*(*t*) = *A_i_a_i_* log(2)2^−*a_i_t*^.

We can therefore rewrite the density in a more convenient form, as the ratio 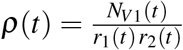. For short time after germination, *r_i_*(*t*) can be safely approximated using a linear function *r_i_* = *B_i_t*, as can be seen in figure 5. In^16^ we have shown that *N*_*V*1_ can be written as *N*_*V*1_(*t*) = *C*2^*ωt*^, with *C* > 0 and *ω* > 0, and we estimated these last parameters. We can therefore derive the following expression for the density, 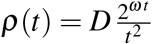, with 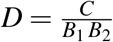. The density diverges for both *t* → 0 and *t* → ∞. In other words, density will show a minimum for an intermediate time, *t*_min_, which defines two distinct growth phases. Note that a minimum always exists if *S*_1_ = *P_n_*(*t*) with *P_n_* a polynomial of order *n* (with the term *n* = 0 to respect the initial condition *S*_1_(0) = 0) or if the time dependence characteristic length generated by the *V*_1_ cloud is of Brownian type (*i.e*. 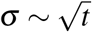). Basically, if the characteristic *S*_1_ area is a *t*-polynomial, two distinct growth phases are identifiable on either side of *t* ≃ 1/*ω*, where *ω* is the characteristic growth parameter of the number of *V*_1_. We can then safely derive an estimation of 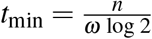, with *n* = 2 in the case where the eigenvalues grow linearly in time.

Let us now compare the empirical density *ρ_o_*(*t*) obtained from the simulation with the theoretical form of *ρ*(*t*) discussed previously. First, we adjusted the spectra of *N*_*V*1_(*t*), *r*_1_(*t*) and *r*_2_(*t*) independently in order to derive *D* and *ω*, allowing to regain *ρ*(*t*). The corresponding density is shown in figure 6A. The grey area corresponds to the theoretical form of *ρ*(*t*) with one standard deviation (see below). The points correspond to the *ρ_o_*(*t*) densities measured individually at each time step from the simulation. The uncertainties for the *ρ_o_*(*t*) points and for the parameters introduced in the theoretical form of *ρ*(*t*) are calculated using the bootstrap method and relying on the Poisson hypothesis for counting process. The general behaviour with a marked minimum is regained in both cases. The empirical minimum (corresponding to *ρ_o_*(*t*)) is found approximately for 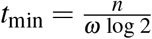, compatible with the value of *t*_min_ calculated by assuming a linear behaviour of the eigenvalues, *t*_min_ ≃ 5 h (corresponding to *ρ*(*t*)).

**Figure 6.**
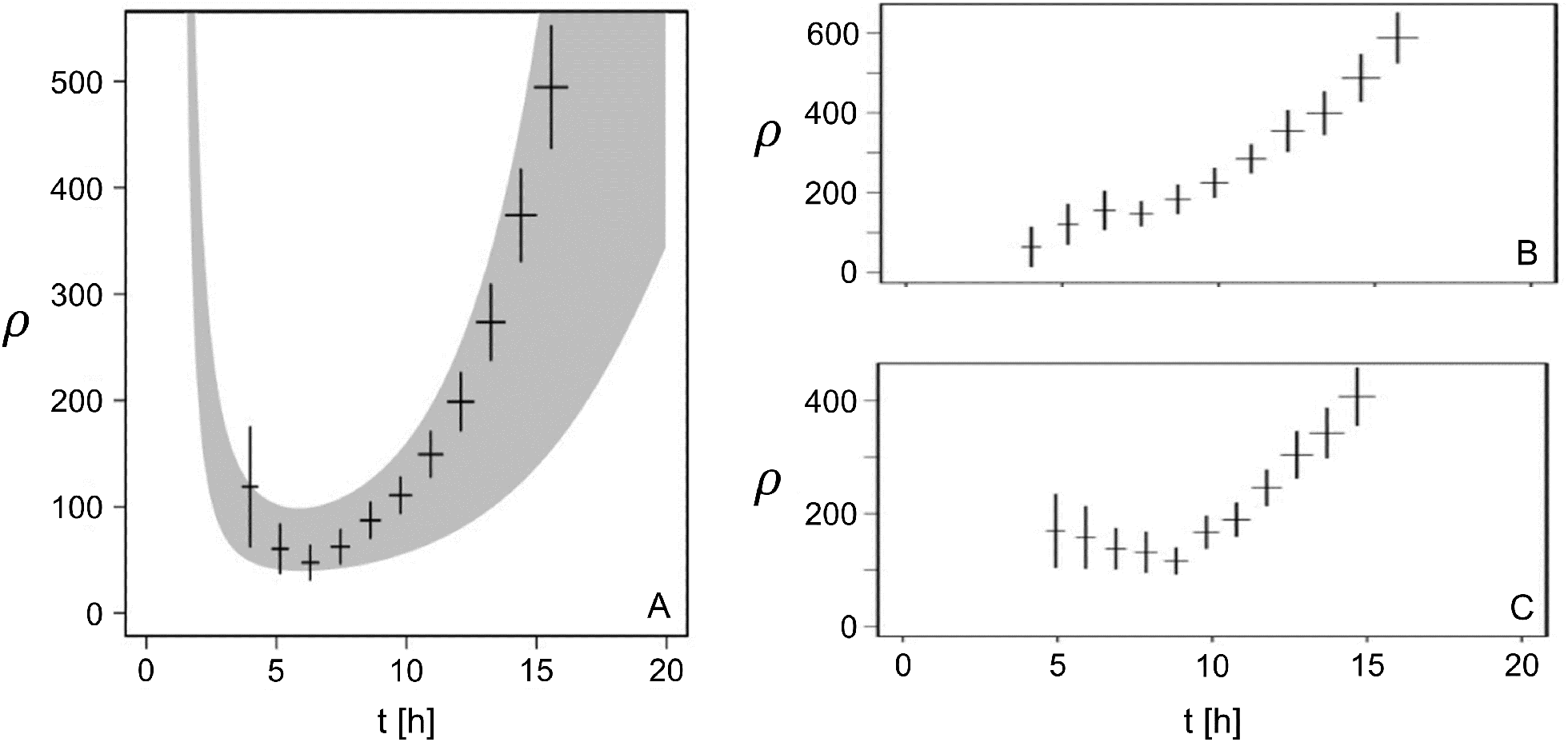
(A) Density versus time for empirical data of simulation (points) and theoretical function 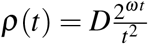 with adjusted parameters (grey area for one standard deviation). (B) same as (A) but with many lateral branches and big angle for operating branch. (C) same as (A) but without lateral branches and small angle for exploratory branch.

We show in figure 6B and C respectively, alternate results from the same analysis method, but with additional constraints. First, we maximized the number of *V*_1_ vertices near the outer ring by prohibiting lateral branching and by setting the angle of the operating branches to the small value of 10 degrees (figure 6B). All material production is then concentrated at the tips of the apexes located favourably in the outer crown. Second, we have chosen parameters that maximize the density at the centre of the network, by increasing the lateral branching frequency and setting the angle of the operating branches to a large value, *i.e*. 120 degrees. The production of new material is then favourably located in the vicinity of the centre (figure 6C). Since the density is written as the ratio of an exponential to a polynomial, we found in both cases that the density grows in the long run. With the chosen parameters, the branching frequency is lower than the reference in the first case and higher in the second. The minimum density being expected for a growth time *t*_min_ which varies in 1/*ω*, we regain that *t*_min_ of the reference is framed by the two proposed variants. We can then distinguish two specific functions for the apical and lateral branches. The former are related to the occupation of the long distance space defining a perimeter, in which the latter will locally densify to exploit the available nutrients.

To conclude, we propose the following prediction regarding the fungi life course. In a homogeneous environment, after germination, the growth of the network presents three distinct life phases, which can be observed through the density.

— During the first phase of growth, which lasts approximately six hours, the growth dynamic maximises the space explored, in order to optimise the colonisation of its distant environment and to favour the future exploitation of available resources. During this extension phase the density is mechanically reduced.
— During the second phase, the lateral branching process appears and it balances the mass distribution in the network, *i.e*. the density remains stable over periods comparable in duration to the first growth phase.
— In a third phase, exploitation of the colonized area becomes the dominant behavior, *i.e*. the capture of resources, and the density must increase significantly. To this end, *P. anserina* can develop lateral branches from any point in its network, which fixes it permanently in its environment. The development of the occupation of the already explored surfaces allows i)to feed the whole network which is far from the apexes and *ii*)to avoid the presence of other competing organisms.

Finally, it is expected that the time at which the density reaches its minimum *t*_min_ depends only on the growth rate *ω* of the thallus. These different growth periods are not specific to the *P. anserina* filamentous fungus network, whose growth can be modeled as a binary tree. For example, different metabolic phases in human life course were reported.^24^ In this case the marker used is the daily energy expenditure, or in intensive form, the energy mass density.

### 2.2 Experimental quantification of the thallus growth

In previous works,^15, 16^ we described the acquisition process and data processing of images of *P. anserina* whole thallus. We obtain from the growth of the thallus a collection of images regularly spaced in time of 18 minutes during the first 20 hours of growth. The spatial resolution of the images (about 1 *μ*m) allows the observation of the fine structure of the hyphae network (roughly 5 *μ*m in diameter). A graph of the network formed by the thallus of *P. anserina* is then reconstructed from each image of the collection, see figure 4. In this graph, the heads (apexes) are degree 1 vertices, and their number is noted A. The network itself is composed of hyphae, degree 2 vertices, whose total length is noted *L*. We used these generic data to calibrate the experimental simulations, as described in section 2.1.1.

In the rest of this article, based on the time series *A* and *L* obtained during the growth of the thallus from an ascospore and over a period of typically 15 hours in a controlled environment, we propose to discuss the temporal dynamics of the density. Following the definition proposed in the previous section 2.1.2, we construct the density at each time step using the eigenvalues of the collection of apex locations and the number of apexes *A*.

#### M2 culture medium

The starting point of each experiment is an ascospore that is placed in standard growing conditions, at a temperature of 27°C, as soon as germination has occurred. We did use of this setup to carry out three complete and independent series of images of the thallus growth, under two standards culture media, M2 and M0. The M2 culture medium is the most commonly used for *P. anserina* growth and reproduction *in vitro*. The carbon source is dextrin, a polysaccharide derived from starch. The M0 culture medium composition is the same as M2, but without a carbon source. It should be noted here that the M0 culture medium does not contain any carbon source but it is known that *P. anserina* is able to partially degrade the cellophane used to maintain the fungus in two dimensions.^14^ However, this carbon source is largely in the minority compared to the standard M2 culture medium. All the protocols including standard culture conditions and media composition for this microorganism can be accessed online, (see^25^ and^14^). Experiments are named respectively M0_*i*_ and M2_*i*_; with *i* = 1,2 or 3 thereafter.

The general base 2 exponential form for a limited growth time of the number of apices *A* = *C*2^*ωt*^ has already been extensively discussed in,^15, 16^ especially the extraction of the parameters *ω* and *C*. The doubling frequency is found in this work respectively at 0.48 ±0.03, 0.46 ±0.02, 0.49 ±0.03 with the culture medium M2 in agreement with the values reported in the table 1.

**Table 1.**
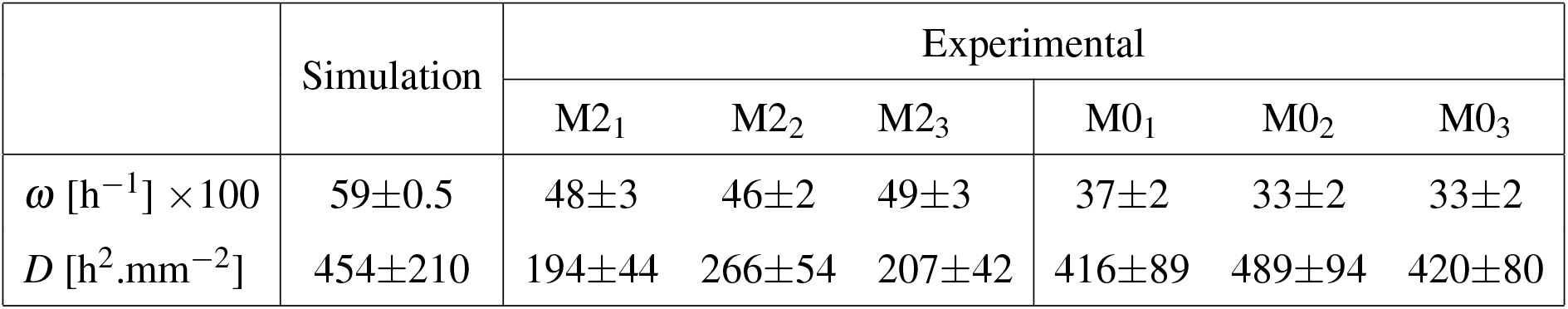
Summary of numerical values for the data from simulation and from the experiment, allowing to regain the density 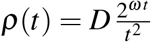. *ω* is the exponential argument of the number of apexes and 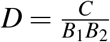 is obtained from the combination of the initial number of apexes *C* and the results of the eigenvalues fitting procedure, see figure 5 and text for details. To ease the reading, *ω* values were multiplied by 100.

|  | Simulation | Experimental |  |  |  |  |  |
| --- | --- | --- | --- | --- | --- | --- | --- |
|  |  | M2<sub>1</sub> | M2<sub>2</sub> | M2<sub>3</sub> | M0<sub>1</sub> | M0<sub>2</sub> | M0<sub>3</sub> |
| $\omega$ [h<sup>-1</sup>] × 100 | 59±0.5 | 48±3 | 46±2 | 49±3 | 37±2 | 33±2 | 33±2 |
| $D$ [h<sup>2</sup>.mm<sup>-2</sup>] | 454±210 | 194±44 | 266±54 | 207±42 | 416±89 | 489±94 | 420±80 |

We will focus in this paper on the evolution of the surface, defined here as the product of the eigenvalues of the cloud formed by the collection of the apex locations at each time step. We have represented in figure 5B the temporal evolution of *r*_1_ and *r*_2_, for the M2_1_ experiment. Both of them clearly follow a linear law, whose slopes *B*_1_ and *B*_2_, about 100 *μ*m.h^−1^, are found to be comparable between them and also compatible with the evolution predicted during the simulation.

This allows us to derive the temporal evolution of the density. We have represented density in figure7A for the experiment M2_1_. We regain the three-step evolution of the density, showing a pronounced minimum. We have also represented *ρ* in superimposition for the same experiment. The behavior for the smallest times is found in the trend but the dynamics is clearly less great. The agreement becomes very good as soon as the growth time exceeds approximately 7 hours, in the vicinity of the density minimum. In order to allow the comparison with the simulation, we extracted *ω* the growth rate and *D_i_* the corresponding prefactor. In accordance with the simulation, we found 0.47±0.03h^−1^ for *ω_i_* and 222 ±50h^2^.mm^−2^ as average value of *D_i_* for the 3 experiments on M2 culture medium. We can also derive an estimation of *t*_min_ using 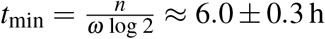, when the eigenvalues are both linear in time (*n* = 2). We have summarized in the table 1 the values extracted from the different experiments and simulations presented.

**Figure 7.**
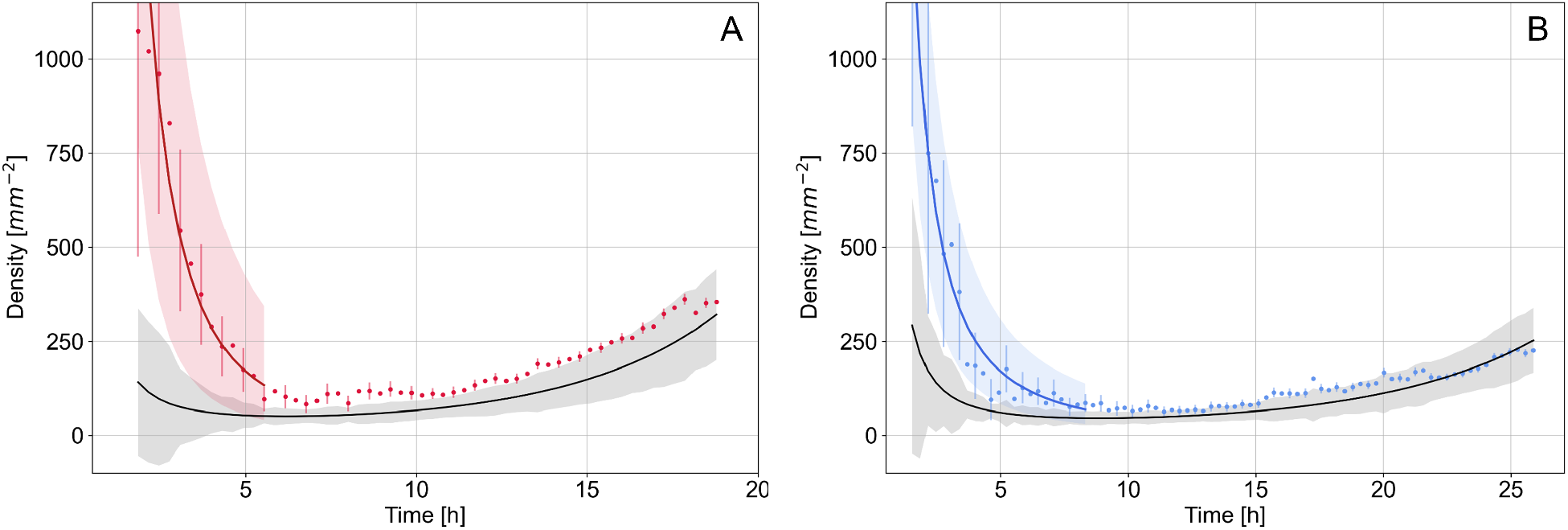
Experimental density in function of time. Culture media are respectively M2_1_ (A) and M0_1_ (B). Circles are density computed at each time step, black solid line is density *ρ* computed from expected laws of apexes number *A* and surface *S* growth. Grey area shows one standard deviation error. Solid red and blue lines are fit based on *β t*^−*α*^. Red and blue areas show one standard deviation. Respectively for M2 and M0 culture media, *α* were found to 2.1±0.3 and 1.6±0.3 with *R*^2^ = 0.96 and 0.95. With *α* = 2 (not shown) we found *R*^2^ =0.96 and 0.76.

#### M0 culture medium

The partial agreement observed is based on growth on M2 culture medium. The question that arises is to discriminate the origin of the dynamics between an effect of the external environment, the available resource, or the stress and a behavior driven by a dynamic inscribed in the fundamental processes of growth and which are thus not expected to be affected either by the resource or by the stress. How do the phases evolve when the metabolism is slowed down by a nutrient-depleted environment? For this purpose we conducted a second triplicate with a nutrient-depleted medium, whose realizations are called M0_*i*_, with *i*=1, 2, 3 in the following.

As expected,^16^ for both *B*_1_ and *B*_2_, the slopes of the eigenvalues, and the growth rate *ω* are reduced by one fourth, respectively to approximately 75 μm.h^−1^ and 0.34±0.02h^−1^ with nutrien-depleted environment. Consequently *D* ≈ 450 ±50 h^2^.mm^−2^ is found to be twice higher than in the M2 condition, see table 1. However, the general dynamics are preserved, leading to density dynamics equivalent to what has been reported previously, with good agreement beyond the first growth phase and a very marked minimum around 10 h.

## 3 Discussion

The various extractions made on the dynamic behavior of the thallus in section 2.1.1 allowed us to feed the modeling of the thallus growth to derive fine predictions. Thus, the density 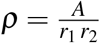, constructed as the ratio of the number of apexes *A* and the product of the eigenvalues (*r*_1_, *r*_2_) of the inertia matrix *I*, consisting of the collection of the apexes locations. The expected behavior is composed of three phases of growth. The first one is a spatial extension, marked by a rapid decrease of the density. The second is a phase of homogeneous extension, where extension and exploitation grow in a density-conserving manner. The third phase shows a dynamic of intense exploitation, where the density increases. A density minimum is found, which depends only on the growth rate ω of the number of apexes A. These phases seem generic and are also observed in human metabolism for example.^24^

We can then compare with direct observations of the growth of *P. anserina*. In both cases, the density dynamics is well found, especially at long time, with a marked minimum, showing a transition between a phase of decrease and a phase of growth. The disagreement with the initial phase of density decay is also found. The times at which the density reaches its minimum on the standard M2 medium, used to calibrate the model, and in the simulation are compatible, found around 6 h in both cases. It is interesting to note that this time is a function only of the polynomial describing the increase of the occupied surface through time, whose order is fixed in this work at *n* = 2, leaving as the only adjustable parameter the growth rate of the apexes number *ω*. Thus, it comes that the nutrient-depleted medium M0, for which *ω* is lower by about a third has mechanically *t*_min_ higher, here found at about 10 h. Thus, the phases of densification are independent of the available resource, which will only have an impact on the speed at which the different phases are explored.

We are now interested in the initial difference between the observed and expected density. Recall that the density is calculated as the ratio of the number of heads *A* to the product of the eigenvalues of the collection of head locations *r*_1_, *r*_2_, which is expected to vary in *t*^−2^. At long time, the behavior of *A* is exponential, *A* ∝ 2^*ωt*^. On the contrary, at short time, when the number of apexes is small, *i.e*. close to unity, the growth does not correspond anymore to the proposed law, it is rather linear, see figure 8A and B. Two processes are responsible for the growth of the number of *A*-heads: apical and lateral branches. The former are specialized^16^ in the exploration of the environment, while the latter are related to the densification of the network. The latter are crucial in the growth of the apexes number. We have seen that the branching statistics follow a random distribution for both types of branching, preceded by a forbidden region, also called apical dominance, see figure 1. However, the numerical values associated with these laws are not identical. The region of apical dominance is of the order of 230 *μ*m, while that of the laterals is found to be of the order of 400 *μ*m. It is this difference that sets the two growth phases. The first lateral branches occur when the initial apical branches exceed the critical length. The velocity *v_h_* of the hyphae from the spore, in the first hours of growth, was found to grow steadily from 0 to 4.5 *μ*m.min^−1^, see figure 8C. Thus, it comes that a new densification phase should appear after about a duration Δ*t* such that 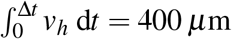, or 7 hours of growth.

**Figure 8.**
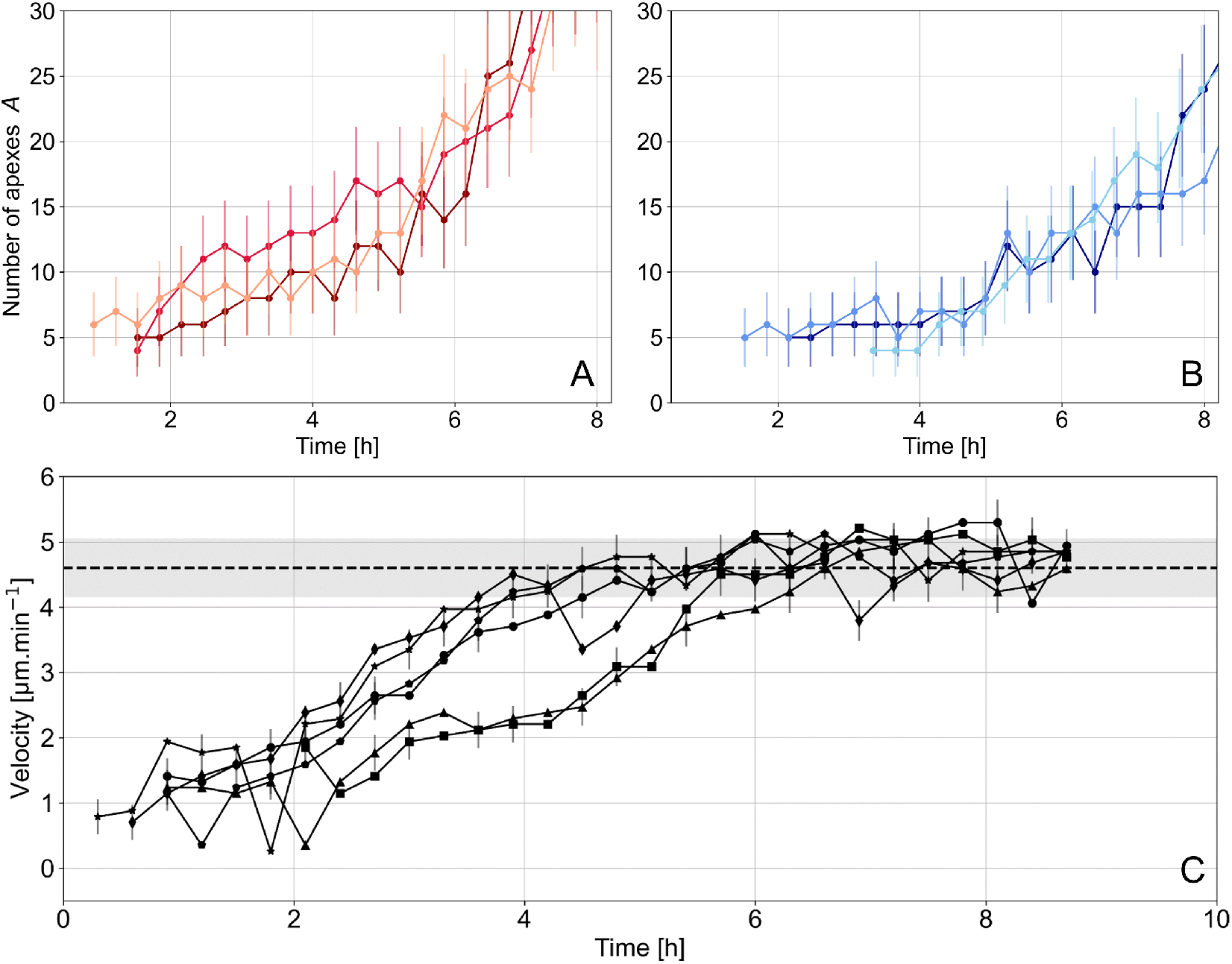
Number of apexes as a function of time for both replicates on M2 medium (A) and M0 medium (B). Only the initial period of growth is shown, defined up to the average time *t*_min_ for the three replicats on M0 medium. (C) Respective velocities of the six initial hyphae. Dashed line corresponds to the average of the asymptotic velocities of 20 apical hyphae (Not shown). We have represented in shaded grey one standard deviation of the distribution of these values

The question that arises then is to know if the initial dynamics depends on the available resource or if it is controlled by the initial conditions (as it can be observed for seeds), *i.e*. the stock of resource present in the ascospore, assumed to be identical in all the experiments carried out. Based on the reasonable assumption of *i*) a linear dependency of the eigenvalues of *I*, *i.e. r*_1_ *r*_2_ ∝ *t*^−2^, and *ii*)*A* is constant in the initial period, then we must find that the density varies in *t*^−2^. We show in figure 7 the fit of the experimental data with the *βt*^−*α*^ law in the time range [0 – 0.9*t*_min_] with *t*_min_ such that the minimum of the density is *ρ*(*t*_min_). It is clear that the proposed fit is much better than the law derived from the long time behavior.

Now let us discuss the numerical values of the *α* exponents found, in relation with the proposed behavioral law. The exponents found are respectively 2.1, 2.5 and 2.3 for M2 and 1.6, 1.9, 1.1 for M0 culture medium, all uncertainties being smaller or equal to 0.3. These exponents are clearly higher in the M2 case, indicating a higher growth rate of *A* the apexes number, but remain in all cases compatible with a slope in *t*^−2^.

Finally, we can test the hypothesis of independence of the dynamics of the initial growth from the available resources. For this, let us set *α* = 2, we then check the dependency of the fit on the boundary conditions. We found that *R*^2^ was in the range 0.6 to 0.9 when the exponent is free. With *α* = 2, it comes that *R*^2^ remain constant or decrease by about 0.1. Given the uncertainties, we cannot conclude that the fit with exponent *α* set to 2 would be better and that the initial growth process is independent of the culture medium. *β* are found for respectively M2 and M0 culture media 2 300 ±200, 5 400 ±400, 2100 ±100 and 3 300 ±400, 4 900 ±400, 11200 ±2 600 h^2^/mm^2^. The values of *β* extracted in this fit are not found to be compatible and allow us to conclude that the first order effect on the initial growth is due to the linear rather than exponential law of the number of apexes *A*.

## 4 Conclusion

In this work we propose the simulation of the growth of the branching network of *P. anserina* based on a binary tree and whose parameters are finely improved from experimental observations. They take into account, in particular, the separation of branching statistics of lateral and apical apexes, the spontaneous curvature of hyphae outside branching events, the average orientation of branches and aging effects due to extreme densification. This allows to construct an expression of the density based on the growth laws of the number of apexes *A* and a surface constructed thanks to the eigenvalues of the matrix of apex locations. A typical behavioral law on densification is then derived, which is expected to follow two phases, separated by a minimum, corresponding to a growth duration whose expression depends only on the growth rate of the apexes and the degree of the polynomial expressing the variation of the occupied surface. The typical behavior is well found experimentally, as well as the dependence of the value of *t*_min_ on the growth rate of *A* and thus on the culture medium. The two phases of densification are explained by the difference in the lengths of the respective apical dominances of the apical and lateral branches. Finally, a deviation from the typical law at the beginning of growth is observed and discussed. It is compatible with a change observed in the growth dynamics of the number of apexes. This one in the initial phase is linear rather than exponential.

## Acknowledgements

We warmly thank Sylvie Cangemi and Aurélien Renault for their expert technical assistance. Clara Ledoux is supported by a PhD scholarship from the Doctoral School MTCI (ED 563). This work is supported by the IdEx Université Paris Cité (ANR-18-IDEX-0001) and by ANR-21-CE40010-01.

## Author contributions

C.L. performed biological experiments, contributed to their analysis and discussed the results. G.R.R. contributed to the data analysis and discussed the results. C.B. discussed the results. Ch.L. contributed to the data analysis and discussed the results. F.C.L., E.H. and P.D. conceived the ideas, performed the experiments and the analysis, interpreted the data and wrote the article. P.D. performed the simulation. All authors reviewed and amended the manuscript.

